# VASP association stabilizes the actin nucleus revealed by molecular dynamics simulations

**DOI:** 10.64898/2026.10.17.759784

**Authors:** Xiaoting Li, Jie Hu, Yu Sun, Congfei Xu, Yiben Fu

**Author notes:** These authors contributed equally to this work (Xiaoting Li, Jie Hu).

## Abstract

Actin nucleation is a rate-limiting step in F-actin assembly. However, early actin trimers are intrinsically unstable, with limited inter-subunit contacts rendering them prone to dissociation. VASP is an important regulator of actin polymerization and assembly, but how it stabilizes unstable trimeric actin nucleus remains poorly understood at the molecular level. Here, we use coarse-grained and all-atom molecular dynamics simulations to characterize the interfacial stability and conformational features of VASP-bound actin trimers. Energetic and interfacial analyses show that VASP preferentially anchors the laterally associated subunit through multivalent interactions and reinforces its interfaces with the longitudinal actin dimer. Within the VASP-bound trimer, the three actin subunits retain distinct local conformational features, as reflected by differences in W-loop gating and SD1-to-SD4 dihedral angle. Across replicate simulations, VASP stabilizes the actin nucleus while preserving subunit-specific conformational heterogeneity, rather than uniformly rigidifying the trimer. These findings provide a molecular basis for VASP-mediated stabilization of early actin nuclei and offer broader mechanistic insights into the regulation of protein assembly.

## Introduction

Actin is a major component of the cytoskeleton, essential in cell shape, migration, adhesion, cytokinesis, and mechanical responses (1). Actin polymerization involves the assembly of globular G-actin monomers into polarized F-actin filaments. Filament formation proceeds through nucleation and subsequent elongation, with nucleation representing a major kinetic barrier in the assembly process (2, 3). During nucleation, two actin monomers first associate to form a longitudinal dimer, followed by the lateral association of a third subunit to generate a trimeric nucleation core (2, 4, 5). However, these early actin assemblies are intrinsically unstable because their inter-subunit contacts are limited, making them prone to dissociation (1, 3, 4). Their lifetime and conformational stability directly influence subsequent filament elongation. Actin nucleation is also accompanied by conformational rearrangements within individual subunits, including local loop dynamics and changes in relative subdomain orientation (4, 6, 7). These conformational changes are closely coupled to the formation and stabilization of inter-subunit interfaces and determine whether the trimer can adopt a nucleation-competent state. Understanding how the unstable actin trimer is stabilized is therefore important for elucidating the molecular regulation of actin polymerization and cytoskeletal remodeling.

As a member of the Ena/VASP family, VASP assembles into tetramers and serves as a key regulator of actin cytoskeletal remodeling (8). Through its EVH2 domain, VASP recruits actin monomers and promotes filament elongation and local actin enrichment, thereby contributing to processes such as filopodia formation and membrane protrusion (9).

Under molecularly crowded conditions, VASP can also undergo liquid–liquid phase separation, which may further modulate actin filament assembly (10). Although the role of VASP in actin filament elongation has been extensively characterized, whether and how VASP contributes to the formation and stabilization of early actin nuclei remains poorly understood at the molecular level (8, 10, 11). In a trimeric actin nucleus, the laterally associated subunit forms relatively limited contacts with the longitudinal actin core and therefore represents a potentially vulnerable component of the assembly (4). Whether VASP preferentially stabilizes this lateral subunit and reinforces its interfaces with the adjacent actin subunits is thus critical to understanding how VASP may promote the formation of a stable actin nucleus.

Actin nucleation is highly transient and dynamic, making it difficult for static structures alone to capture the interfacial rearrangements and conformational fluctuations associated with their stabilization (2–4) . Molecular dynamics simulations provide a complementary approach for resolving these processes over time. Coarse-grained simulations enable extended sampling of global conformational evolution, relative subunit motions, and large-scale interface rearrangements, whereas all-atom simulations provide the resolution required to characterize residue-level interactions and local structural dynamics (12, 13). Combining long-timescale coarse-grained sampling with all-atom backmapping and refinement therefore provides a multiscale framework for linking global conformational evolution to the molecular interactions underlying actin-trimer stabilization.

In this study, we constructed a VASP-bound actin trimer and combined coarse-grained molecular dynamics with all-atom simulations initiated from backmapped structures to investigate how VASP stabilizes early actin nucleation. We examined the VASP–actin interface, the internal interfaces within the actin trimer, and subunit-specific conformational features, including W-loop gating and SD1-to-SD4 orientation. Our repeated simulations show that VASP anchors the laterally associated actin subunit and reinforces its interactions with the longitudinal dimer while preserving distinct local conformational states among the three actin subunits. These results suggest a molecular mechanism by which VASP stabilizes a nucleation-competent actin trimer during the early stages of actin assembly.

## Results

### VASP remains stably associated with the actin trimer during coarse-grained simulations

We first examined the long-timescale structural evolution of the actin–VASP complex using Martini 3 coarse-grained molecular dynamics simulations (12). The actin trimer consists of three subunits, A1, A2, and A3, with A2 occupying the laterally associated position relative to the longitudinal A1/A3 dimer (4) **(Fig. 1A)**, whereas VASP adopts a tetrameric assembly (9, 10) **(Fig. 1B)**. Throughout the 3 μs simulations, VASP remained associated with the actin trimer, with no global dissociation of the complex. Rather than being confined to a single localized interface, multiple regions of VASP simultaneously contacted A1, A2, and A3, resulting in an extensive multivalent interface across the actin trimer **(Fig 1C)**.

**Figure 1.**
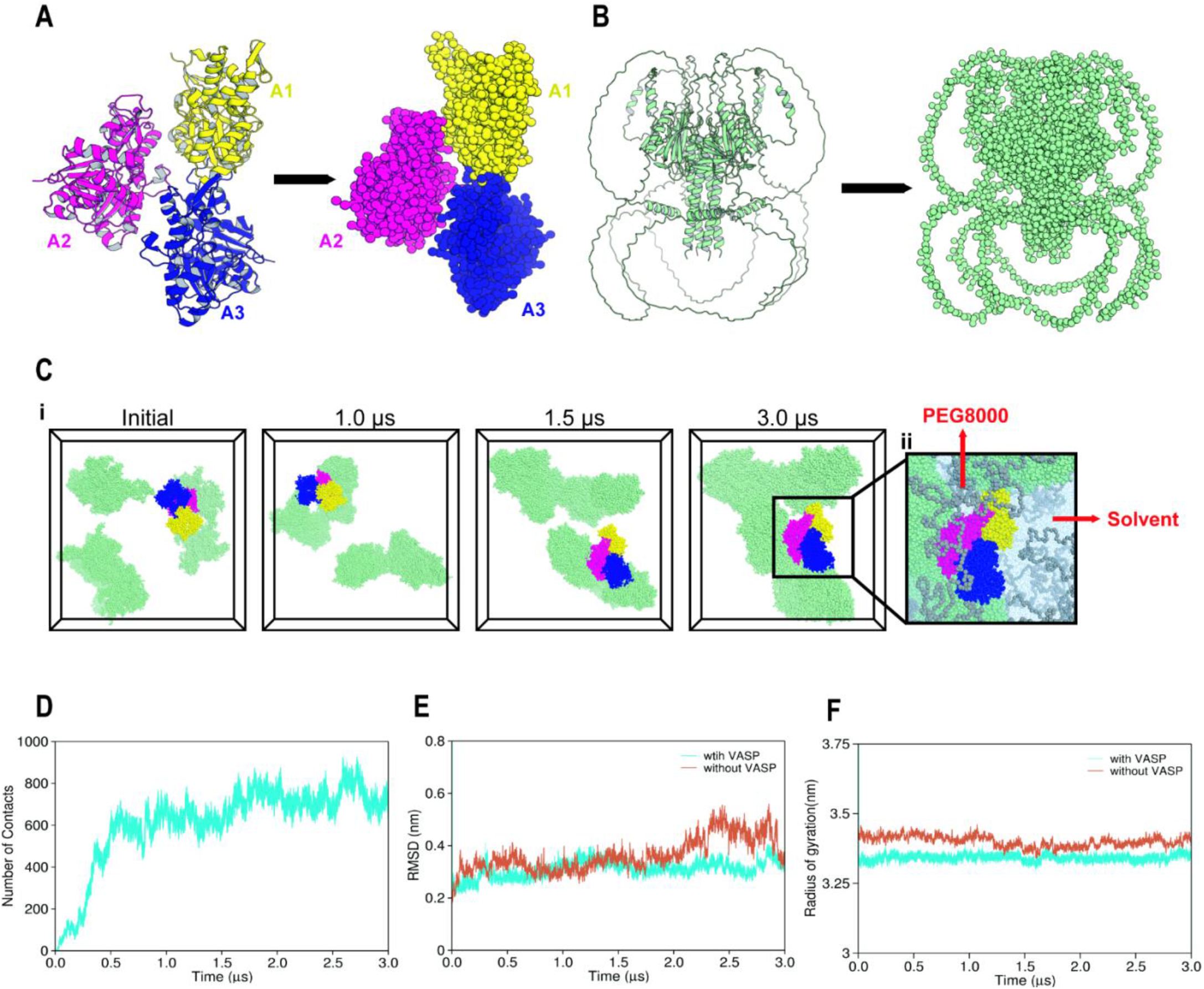
Stability of the Actin–VASP complex in coarse-grained simulations. **(A)** All-atom and Martini 3 coarse-grained representations of the actin trimer. Subunits A1, A2, and A3 are shown in yellow, magenta, and blue, respectively. **(B)** All-atom and Martini 3 coarse-grained representations of the VASP tetramer, shown in green. **(C)** Representative snapshots of the Actin–VASP–PEG8000 coarse-grained system at 0, 1, 1.5, and 3 μs. The enlarged view shows the actin– VASP complex at the end of the simulation, with PEG8000 and solvent shown in the background. **(D)** Time evolution of the number of actin–VASP contacts during the 3 μs coarse-grained simulation. **(E)** Root mean square deviation (RMSD) of the actin trimer in the actin–VASP and actin-only systems. **(F)** Radius of gyration (Rg) of the actin trimer in the actin–VASP and actin-only systems.

The persistence of the actin–VASP association was first evaluated by monitoring interprotein contacts throughout the coarse-grained simulations. Actin–VASP contacts remained abundant over the 3 μs trajectories, with no sustained decrease that would indicate progressive dissociation. Consistently, representative late-stage conformations showed that VASP remained closely associated with the actin trimer, and maintained contacts with multiple actin subunits **(Fig. 1D)**. These observations indicate that the initially assembled actin–VASP complex remained stably associated over the simulated timescale. We next examined whether VASP binding affected the global structural stability of the actin trimer. With VASP, the root mean square deviation (RMSD) of actin trimer reached a relatively stable range after the initial structural relaxation and showed no progressive increase during the later stages of the simulations. The radius of gyration (Rg) similarly remained within a relatively narrow range, indicating preservation of the overall compactness of the trimer. Without VASP, the actin trimer exhibited larger late-stage RMSD deviations and greater variations in Rg, consistent with increased global structural variability **(Fig. 1E and F)**. Together, these results show that VASP remains stably associated with the actin trimer through persistent multivalent contacts and that the presence of VASP is accompanied by reduced global structural variation of the trimer over the simulated timescale.

### VASP preferentially interacts with the lateral A2 subunit

We next examined how VASP interactions were distributed among the three actin subunits. Buried surface area (BSA) analysis revealed that the A2–VASP interface was substantially larger than either the A1–VASP or A3–VASP interface **(Fig. 2A)**, indicating that the laterally associated A2 subunit form the most extensive contact with VASP. Consistent with this interfacial asymmetry, MM/GBSA analysis yielded a more favorable interaction energy for A2–VASP than for A1–VASP or A3–VASP **(Fig. 2B)**. Together, these analyses identify A2 as the principal VASP-interacting subunit within the actin trimer.

**Figure 2.**
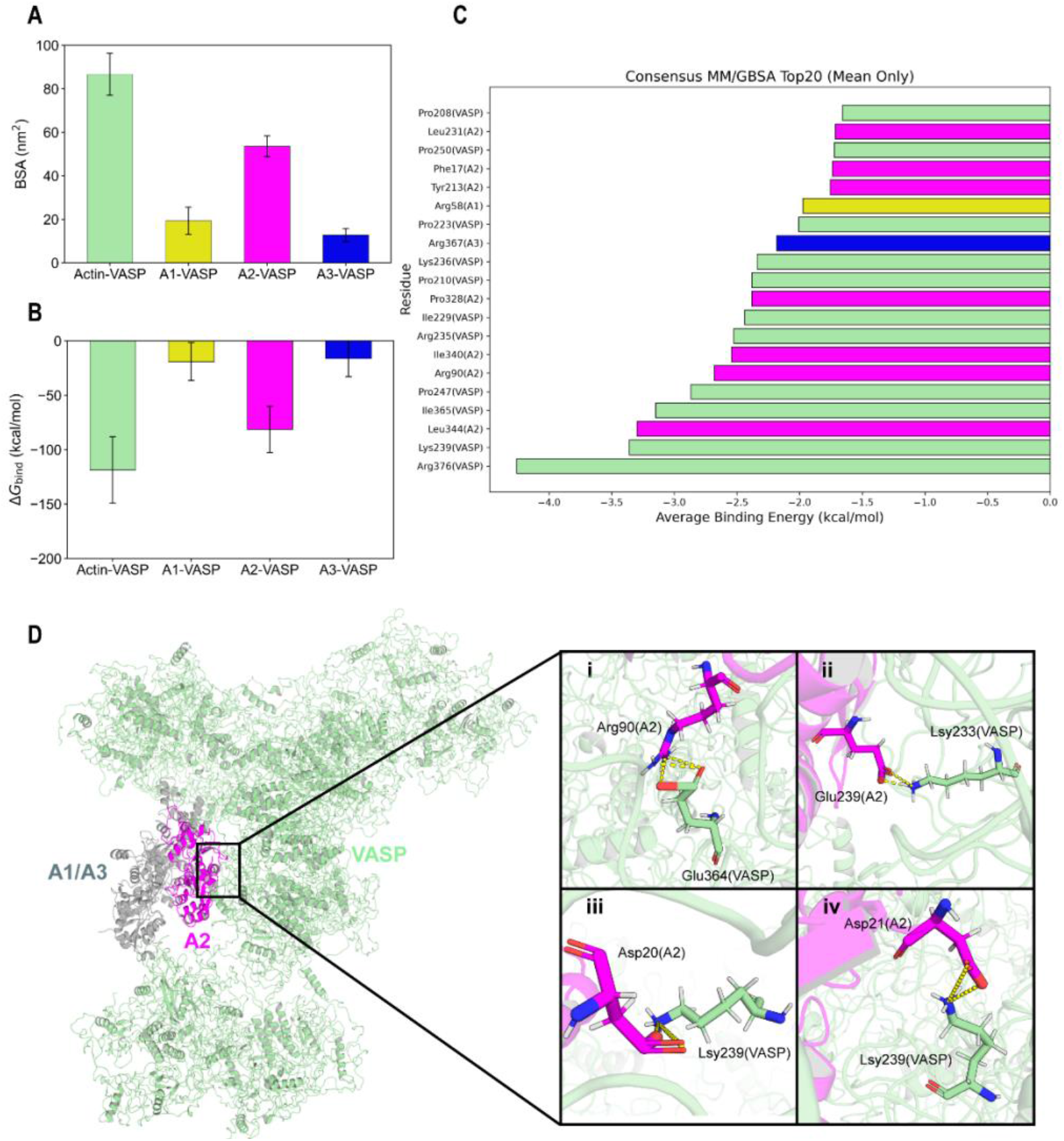
VASP preferentially interacts with the lateral A2 subunit of the actin trimer. **(A)** Buried surface areas (BSAs) of the overall actin–VASP interface and the A1–VASP, A2–VASP, and A3–VASP interfaces. **(B)** MM/GBSA binding energies of the corresponding interfaces. **(C)** Per-residue decomposition of the actin– VASP binding energy showing the 20 residues with the largest energetic contributions. Colors indicate the corresponding actin subunit or VASP component. **(D)** Representative structure of the actin–VASP complex. A2 is shown in magenta, A1 and A3 in gray, and VASP in green. The dashed box indicates the A2–VASP interface. Close-up views of representative high-occupancy interactions at the A2–VASP interface. Panels i to iv show Arg90(A2)–Glu364(VASP), Glu329(A2)– Lys233(VASP), Asp20(A2)–Lys239(VASP), and Asp21(A2)–Lys239(VASP), respectively. Key residues are shown as sticks, and dashed lines indicate hydrogen-bond or salt-bridge interactions.

Per-residue energy decomposition further resolved the molecular determinants of the Actin–VASP interface. Among the 20 residues with the most favorable energetic contributions, multiple residues originated from A2, including Phe17, Leu231, Tyr213, Pro328, Arg90, Ile340, and Leu344, together with contributing residues from VASP **(Fig. 2C)**. The distribution of energetic contributions across multiple residues on both interaction partners indicates that the A2–VASP interface is supported by a broad interaction network rather than by a single dominant contact (14).

Hydrogen-bond and salt-bridge analyses further identified persistent polar and electrostatic interactions at the A2–VASP interface. High-occupancy contacts were predominantly concentrated between A2 and VASP, with prominent residue pairs including Asp20(A2)–Lys239(VASP), Asp21(A2)–Lys239(VASP), Glu329(A2)–Lys233(VASP), Arg90(A2)– Glu364(VASP), and Glu236(A2)–Lys233(VASP) **(Fig. 2D)**. Notably, Asp20(A2)–Lys239(VASP), Asp21(A2)– Lys239(VASP), and Glu329(A2)–Lys233(VASP) were identified in both the hydrogen-bond and salt-bridge analyses (15), indicating that these residue pairs form persistent polar and electrostatic contacts within the A2–VASP interface (16–18).

Together, these energetic and residue-level analyses demonstrate that VASP preferentially engages the lateral A2 subunit through an extensive multiresidue interaction network. This preferential A2 interaction provides a structural basis for examining how VASP association influences the internal interfaces of the actin trimer.

### VASP reinforces A2-associated internal interfaces within the actin trimer

Following the preferential association of VASP with the lateral A2 subunit, we next examined whether VASP association was accompanied by changes in the internal actin–actin interfaces. We first monitored the buried surface areas (BSAs) of the A1–A3, A2–A1, and A2–A3 interfaces throughout the 3 μs coarse-grained simulations to assess their long-timescale evolution with and without VASP. The A1–A3 interface displayed broadly similar BSA profiles under the two conditions, with only modest differences throughout the simulations **(Fig. 3A)**, indicating that this longitudinal interface was comparatively insensitive to VASP association. By contrast, more pronounced differences were observed at the A2-associated interfaces. The A2–A1 interface maintained a larger buried surface area and exhibited smaller fluctuations with VASP than without VASP **(Fig. 3A)**. An even greater difference was observed for A2–A3: with VASP, this interface retained a substantial buried surface area during the later stages of the simulation, whereas its BSA progressively decreased in the absence of VASP **(Fig. 3A)**. These observations indicate that VASP association preferentially preserves the internal interfaces involving the lateral A2 subunit. Interpretation of the BSA changes in the absence of VASP requires consideration of the accompanying global structural rearrangement. As indicated by the larger RMSD deviations and Rg fluctuations **(Fig. 1E and F)**, the actin trimer underwent greater structural reorganization during the later stages of the simulation without VASP. Changes in BSA under this condition therefore do not necessarily represent simple strengthening or weakening of an otherwise unchanged interface, but may also reflect inter-subunit sliding and rearrangement. By contrast, the trimer retained larger A2-associated buried interfaces with VASP while exhibiting comparatively smaller global structural variations. These results suggest that preferential VASP association with A2 helps preserve the contacts between the lateral subunit and the neighboring A1/A3 longitudinal core.

**Figure 3.**
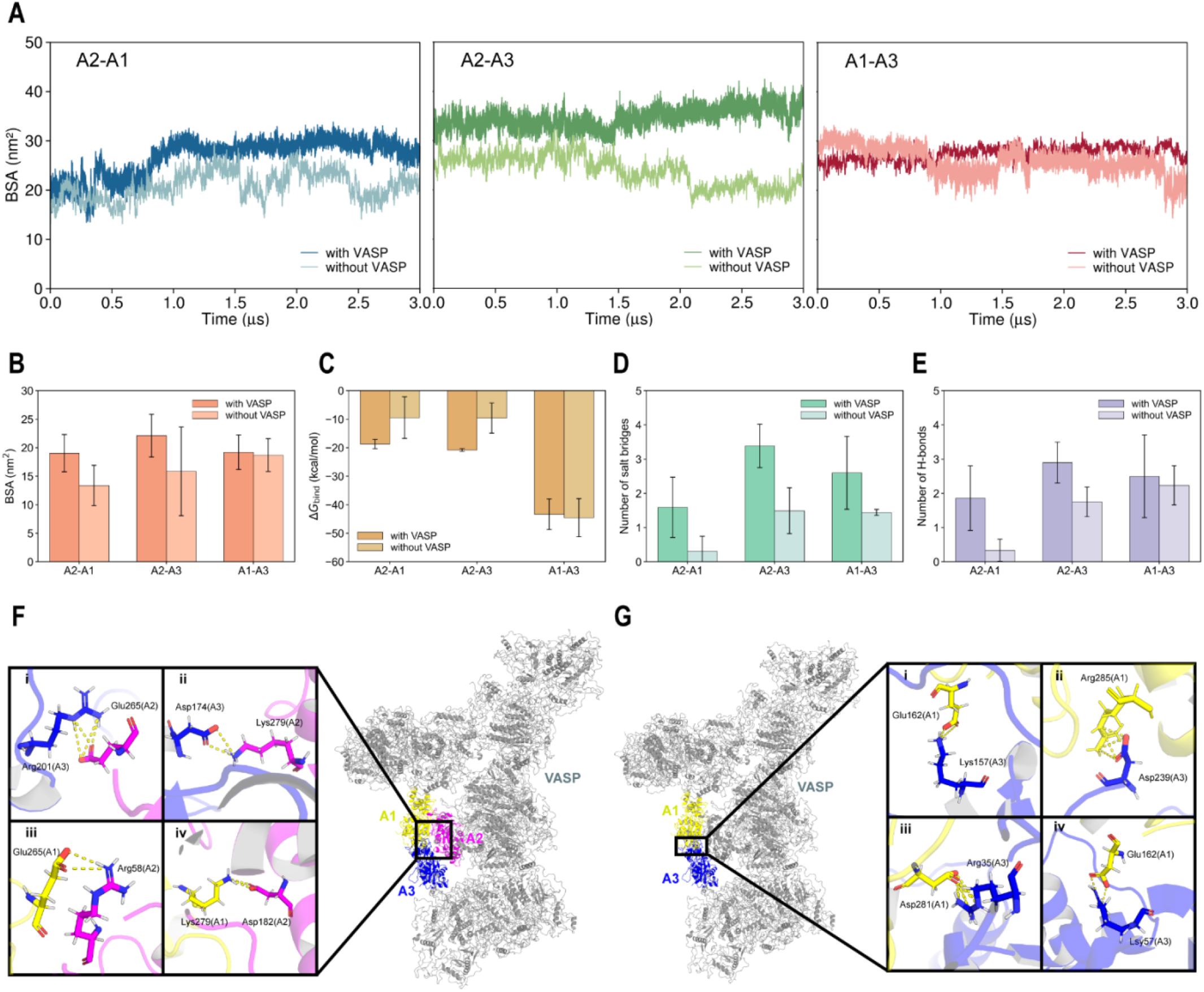
VASP preferentially preserves A2-associated internal interfaces within the actin trimer. **(A)** Time evolution of the buried solvent-accessible surface area (BSA) of the A2–A1 (left), A2–A3 (middle), and A1–A3 (right) interfaces during the 3 μs coarse-grained simulations in the presence and absence of VASP. **(B)** Comparison of the BSAs of the A2–A1, A2– A3, and A1–A3 interfaces over the final 20 ns of the all-atom simulations. **(C)** Comparison of the MM/GBSA binding energies of the three internal actin–actin interfaces over the final 20 ns of the all-atom simulations. **(D)** Numbers of salt bridges at the three internal actin–actin interfaces. **(E)** Numbers of hydrogen bonds at the three internal actin–actin interfaces. **(F)** Representative structure and close-up views of the A2–A1/A3 interfaces, highlighting representative hydrogen-bond and salt-bridge interactions between A2 and the neighboring A1/A3 core. **(G)** Representative structure and close-up views of the A1– A3 interface. Key residues are shown as sticks, and dashed lines indicate representative hydrogen-bond or salt-bridge interactions.

To determine whether these interface-specific differences were retained at atomic resolution, we next compared the BSAs of the three internal actin–actin interfaces over the final 20 ns of the all-atom trajectories. Consistent with the coarse-grained simulations, the A2–A1 and A2–A3 interfaces exhibited larger average BSAs with VASP than without VASP, with the most pronounced difference observed for A2–A3. By contrast, the A1–A3 interface showed only a modest difference between the two conditions **(Fig. 3B)**. Thus, both coarse-grained and all-atom BSA analyses indicate that the influence of VASP on trimer organization is preferentially concentrated at the A2-associated interfaces rather than uniformly distributed across all three internal interfaces.

We next evaluated the energetic characteristics of these interfaces using MM/GBSA analysis over the final 20 ns of the all-atom trajectories. Among the three interfaces, A1–A3 exhibited the most favorable MM/GBSA energy under both conditions, consistent with its persistent longitudinal association. VASP affected the three interfaces differently: the A2–A1 and A2–A3 interactions became energetically more favorable in the presence of VASP, whereas the A1–A3 interface showed only a modest energetic difference **(Fig. 3C)**. These energetic trends were consistent with the BSA results and further support a preferential effect of VASP on the A2-associated internal interfaces.

To further characterize the molecular interactions underlying these interfacial differences, we quantified hydrogen bonds and salt bridges at each internal actin–actin interface. The A2–A1 and A2–A3 interfaces exhibited more hydrogen bonds with VASP than without VASP, whereas the A1–A3 interface showed only a modest difference **(Fig. 3E)**. Salt-bridge analysis revealed a similar overall trend. Although salt-bridge numbers increased at all three interfaces in the presence of VASP, the smallest difference was observed at A1–A3 **(Fig. 3D)**. Together, these analyses indicate that VASP association is accompanied by increased polar and electrostatic contacts within the A2-associated interfaces, whereas the pre-existing longitudinal A1–A3 interface is comparatively less affected. Representative structural views further illustrated the distinct interaction patterns at these interfaces. At the A2-associated interfaces, multiple polar and electrostatic contacts connected A2 with both A1 and A3 **(Fig. 3F)**. In comparison, the A1–A3 longitudinal interface retained a distinct interaction network that was less affected by VASP association **(Fig. 3G)**. These representative interactions are consistent with the interface-selective trends observed in the BSA, MM/GBSA, hydrogen-bond, and salt-bridge analyses.

Taken together, the long-timescale coarse-grained BSA analysis and the all-atom BSA, MM/GBSA, hydrogen-bond, and salt-bridge analyses consistently support a model in which preferential VASP anchoring of the lateral A2 subunit is coupled to reinforcement of its internal contacts with the neighboring A1/A3 core. This selective stabilization of the A2-associated interfaces provides a molecular basis for the enhanced persistence of the actin trimer in the presence of VASP.

### VASP modulates subunit-specific conformational states within the actin trimer

Having established that VASP preferentially anchors A2 and reinforces the A2-associated internal interfaces, we next examined whether this interfacial stabilization was accompanied by changes in the conformational states of the actin trimer. We characterized these changes at three structural levels: local W-loop gating, global subdomain twisting, and positioning of the SD4-associated region. The W-loop gate distance, SD1–SD4 dihedral angle, and Asp239 displacement were used as corresponding conformational indicators (4, 5, 11, 18) **(Fig. 4)**.

**Figure 4.**
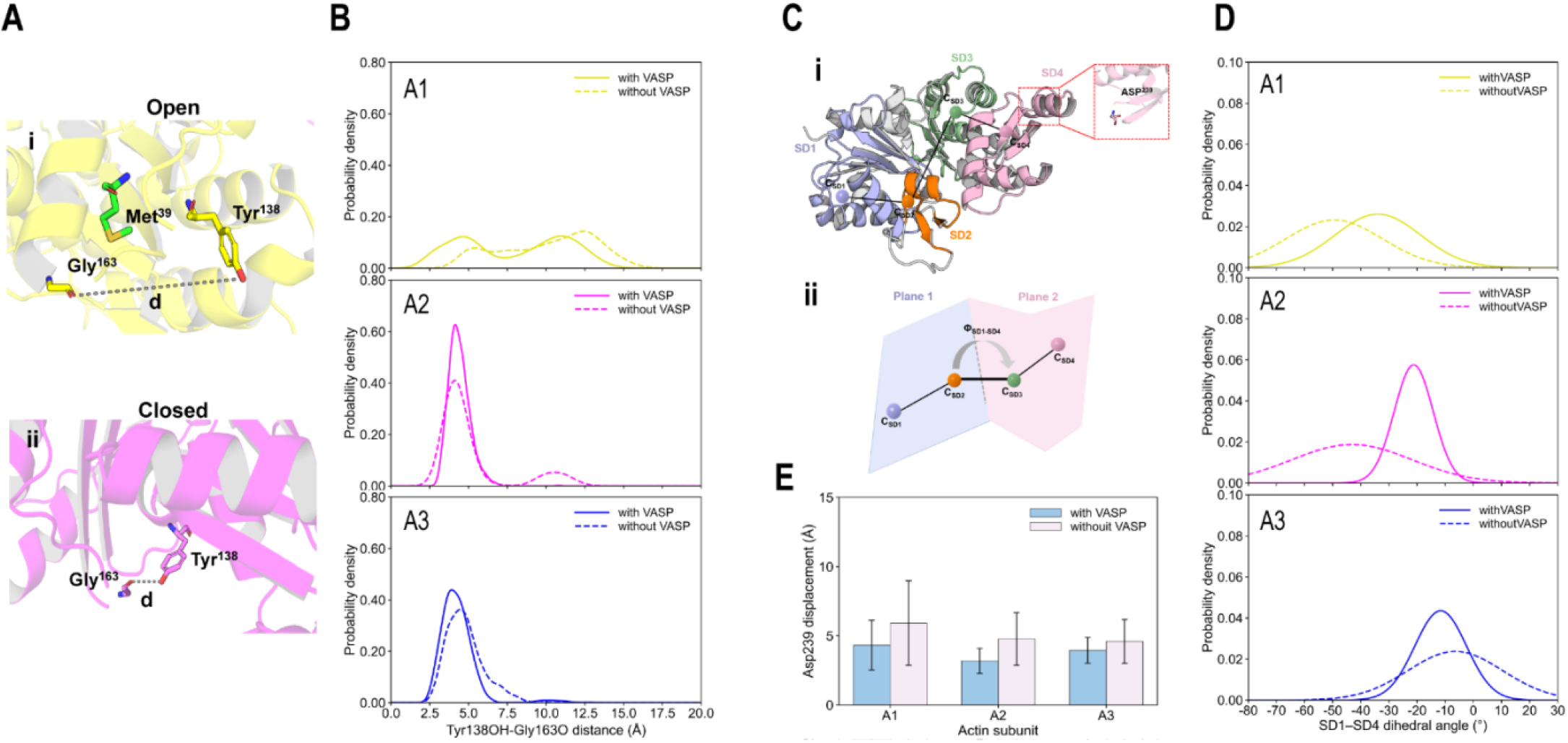
VASP differentially affects conformational states within the actin trimer. **(A)** Schematic representation of the W-loop gate. Tyr138 and Gly163 are shown as sticks, and the dashed line indicates the distance between the Tyr138 side-chain hydroxyl oxygen and the Gly163 backbone carbonyl oxygen. **(B)** Probability density distributions of the W-loop gate distance in A1, A2, and A3, respectively, in the presence and absence of VASP. Solid and dashed lines represent the actin–VASP and actin-only systems, respectively. **(C)** Schematic definition of the SD1–SD4 dihedral angle. The four actin subdomains are shown in different colors, and the centers of SD1, SD2, SD3, and SD4 (C_SD1_, C_SD2_, C_SD3_, and C_SD4_) define the SD1–SD4 dihedral angle, Φ_SD1–SD4_. The position of Asp239 in the SD4-associated region is shown as sticks and labeled. **(E)** Asp239 displacement in A1, A2, and A3 in the presence and absence of VASP. Blue and light pink bars represent the actin–VASP and actin-only systems, respectively. Data are shown as mean ± SD. Lower displacement values indicate closer positional correspondence of Asp239 to the F-actin reference geometry.

First, we examined the conformational dynamics of the W-loop gate. The W-loop controls access to the hydrophobic pocket that accommodates Met39 from the D-loop of a neighboring actin subunit, and therefore represents a local structural feature relevant to inter-subunit docking during actin assembly (4, 19). The gate state was characterized by the distance *d* between the side-chain hydroxyl oxygen of Tyr138 and the backbone carbonyl oxygen of Gly163 within each actin subunit. A shorter distance corresponds to closure of the W-loop gate, whereas a larger distance reflects gate opening (4). In the closed state, hydrogen bonding across the gate restricts access to the Met39-binding pocket, whereas gate opening exposes this site. We analyzed this parameter separately for A1, A2, and A3. The distance distributions in the presence and absence of VASP were compared to assess subunit-specific effects of VASP on W-loop dynamics **(Fig. 4B)**.The W-loop distance exhibited pronounced fluctuations and a bimodal distribution, corresponding to two distinguishable gating states. Based on the separation between the two populations, conformations with *d* < 7.5 Å were classified as closed, whereas those with *d* ≥ 7.5 Å were classified as open. The distributions differed substantially among the three actin subunits. In the absence of VASP, A1 exhibited a broad distribution spanning both low- and high-distance populations, indicating substantial sampling of both closed and open states. By contrast, A2 and A3 were predominantly distributed around 4 to 5 Å and therefore preferentially sampled the closed state.VASP association altered the W-loop distributions in a subunit-dependent manner. In the presence of VASP, the distributions of A1 and A3 shifted toward shorter distances while retaining their overall distribution patterns, whereas A2 exhibited the most pronounced response. In the absence of VASP, A2 retained a detectable open-state population; with VASP, its distribution shifted almost entirely toward the closed state. The VASP-induced shift of A2 toward a more persistently closed W-loop gate indicates that VASP binding alters the local gating dynamics of the Met39-binding pocket. Despite these changes, distinct W-loop distributions were retained among A1, A2, and A3, indicating that VASP does not eliminate subunit-specific local conformational heterogeneity within the actin trimer.

We next used the SD1–SD4 dihedral angle to characterize the global twisting state of each actin subunit. This coordinate describes the relative orientation of the four actin subdomains and provides a structural measure distinguishing the more twisted G-actin-like state from the flatter F-actin-like state (6). In reference structures, the SD1–SD4 dihedral angle is approximately −23.6° for G-actin, whereas F-actin adopts a substantially flatter value of approximately −6.2°. Previous structural analysis further showed that different subunits within a trimeric actin nucleus can occupy distinct states along this conformational coordinate (4). We therefore compared the SD1–SD4 dihedral-angle distributions of A1, A2, and A3 in the presence and absence of VASP **(Fig. 4C and D)**. In the absence of VASP, all three actin subunits exhibited distinct SD1– SD4 dihedral-angle distributions, indicating substantial conformational heterogeneity within the trimer. A2 showed the narrowest distribution, suggesting relatively restricted fluctuations along this twisting coordinate, whereas A3 displayed a dominant population centered near −10°, close to the flattened F-actin-like range. VASP association affected the three subunits differently **(Fig. 4D)**. The peak of A1 was shfited to a larger value, but the distribution remained largely unchanged. A3 retained a similar peak position but exhibited a markedly narrower distribution, indicating reduced fluctuations in its global twisting state. The most pronounced change was observed for A2. In the absence of VASP, the A2 distribution was centered near −55°, substantially more negative than the canonical G-actin reference value. In the presence of VASP, this distribution shifted toward approximately −25°, bringing A2 substantially closer to the canonical G-actin-like twisting range. At the same time, the distribution became broader, indicating increased conformational sampling along the SD1–SD4 twisting coordinate. Thus, VASP does not simply drive all three subunits toward a common filament-like conformation, but instead modulates their conformational states in a strongly subunit-dependent manner, with A2 showing the largest response.

We further examined Asp239 displacement as a positional indicator of the SD4-associated region relative to the F-actin reference geometry (**Fig. 4C)**. For each subunit, the displacement of Asp239 from its corresponding position in the barbed-end F-actin reference structure (PDB ID: 8F8R) was quantified following structural alignment (20). Previous structural analyses similarly used the corresponding SD4 residue as a positional marker for deviations from canonical filament geometry (4). Asp239 displacement was generally reduced for A1, A2, and A3 in the presence of VASP, indicating closer positional correspondence of this SD4-associated region to the F-actin reference geometry (4)**(Fig. 4C and E)**. Together with the SD1– SD4 dihedral analysis, these results show that VASP modulates actin conformation in a subunit-specific manner rather than uniformly rigidifying the trimer.

Together, the W-loop gate distance, SD1–SD4 dihedral angle, and Asp239 displacement analyses demonstrate that VASP affects actin conformation across multiple structural levels, including local gating dynamics, global subdomain twisting, and SD4-associated structural positioning. Importantly, these effects are strongly subunit dependent: A1, A2, and A3 exhibit distinct conformational responses to VASP association. Thus, rather than uniformly rigidifying or homogenizing the actin trimer, VASP selectively reshapes its conformational landscape while preserving subunit-specific structural heterogeneity.

## Method

### Multiscale system construction and molecular dynamics simulations

To capture the long-timescale dynamics of the Actin–VASP complex while retaining atomic-level resolution for interfacial analysis, we combined Martini 3 coarse-grained (CG) and all-atom (AA) molecular dynamics simulations. CG models were first constructed from the initial molecular structures and used to sample the long-timescale structural dynamics of the Actin–VASP system. Representative equilibrated configurations were subsequently backmapped to all-atom structures using CG2AT. The backmapped systems were further subjected to all-atom molecular dynamics simulations to relax local structural distortions introduced during backmapping and to enable quantitative analyses of binding energetics, interfacial interactions, and conformational dynamics.

A coarse-grained system comprising an actin trimer, a VASP tetramer, and PEG8000 was constructed to investigate VASP-mediated stabilization of the actin trimer during nucleation. PEG8000 was included as a macromolecular crowding agent to mimic a crowded condensate-like environment, because previous studies have shown that PEG8000 promotes VASP condensate formation and supports VASP-mediated actin polymerization and bundling under crowded conditions(10, 21). The initial actin trimer structure was obtained from the crystallographic structure (PDB ID: 9JUS) (4), and missing loop regions were modeled using MODELLER (22). The three actin subunits were designated A1, A2, and A3, with A2 corresponding to the laterally associated subunit and A1/A3 forming the longitudinal dimer. The actin trimer was coarse-grained using the Martini 3 force field (12), with an elastic network applied to preserve its overall folded conformation. Because no experimentally determined structure of the full-length VASP tetramer is currently available, its structure was predicted using AlphaFold 3 (23), with each monomer comprising 380 amino acid residues. VASP contains both structured and intrinsically disordered regions. Accordingly, Martini 3 IDP-related parameters were applied to the disordered segments (24), whereas elastic-network restraints were introduced for the structured regions (25). The predicted VASP tetramerization domain showed close structural agreement with the experimentally determined tetrameric structure (PDB ID: 1USE), with an overall tetrameric Cα RMSD of 1.71 Å (Figure S1). During the subsequent all-atom MD simulation of the full-length VASP tetramer, the tetramerization core remained structurally stable, with its Cα RMSD fluctuating predominantly around 0.2–0.25 nm. No spontaneous dissociation of the tetrameric assembly was observed during the simulation. Together, these observations support the structural plausibility of the predicted tetramerization architecture and the suitability of the AlphaFold 3 model as the starting structure for the present simulations. Coarse-grained PEG8000 molecules were generated using polyply (26), with each polymer chain containing 182 polyethylene oxide (PEO) repeat units. The actin trimer, VASP tetramer, and PEG8000 molecules were subsequently assembled in the simulation box using Packmol to generate the complete coarse-grained actin– VASP–PEG system (27).

Coarse-grained molecular dynamics simulations were performed using GROMACS 2024.6 (28). Each system was energy-minimized using the steepest descent algorithm for up to 50,000 steps, followed by sequential equilibration consisting of 1 ns in the NVT ensemble, 5 ns of initial NPT equilibration, and a further 20 ns of NPT equilibration. A time step of 10 fs was used during NVT equilibration and 20 fs during subsequent NPT equilibration and production simulations. The temperature was maintained at 310 K using the velocity-rescaling (v-rescale) thermostat (29). Pressure was maintained at 1 bar using isotropic coupling, with the C-rescale barostat applied during the initial NPT equilibration and the Parrinello– Rahman barostat during the subsequent NPT equilibration and production simulations (30). Electrostatic interactions were treated using the reaction-field method with a relative dielectric constant of 15. Coulomb and van der Waals interactions were truncated at 1.2 nm, with the van der Waals potential shifted smoothly to zero using the potential-shift-Verlet scheme. Three independent 3-μs production simulations were performed for each coarse-grained system. Reproducibility among independent trajectories was assessed using the root mean square deviation (RMSD) and radius of gyration (Rg) of the actin trimer, together with the persistence of actin–VASP contacts. Representative equilibrated configurations from the late stages of the coarse-grained trajectories were subsequently selected for backmapping.

Selected coarse-grained configurations were backmapped to all-atom structures using CG2AT version 2023.6.30 (13). Before backmapping, PEG8000 and other components not required for subsequent protein-interface analyses were removed, retaining the complete actin trimer and VASP tetramer (31). Backmapping was performed using the CHARMM36 force field (July 2020 release) (32). The resulting all-atom structures were subjected to initial structural relaxation and placed in simulation boxes with a minimum protein-to-box distance of 1.2 nm. Each system was solvated with TIP3P water, neutralized with counterions, and adjusted to a final NaCl concentration of 0.15 M.

All-atom molecular dynamics simulations were performed using GROMACS 2024.6. Each solvated system was energy-minimized using the steepest-descent algorithm for up to 50,000 steps, followed by 100 ps of NVT equilibration and 100 ps of NPT equilibration. Production simulations were subsequently performed for 120 ns using a 2-fs time step, with at least three independent simulations conducted for each selected backmapped configuration. Following backmapping from the equilibrated CG trajectories, the actin trimer underwent an initial conformational adjustment during the all-atom simulations before reaching a relatively stable RMSD plateau. Across all three independent simulations, the RMSD stabilized after approximately 60 ns, suggesting that the systems had reached a structurally stable regime. These results indicate that the 120 ns simulation duration was sufficient to capture the conformational equilibration and provide a reliable trajectory window for subsequent structural analyses (Figure S2).Covalent bonds involving hydrogen atoms were constrained using the LINCS algorithm (33), and long-range electrostatic interactions were treated using the particle mesh Ewald (PME) method (25). A Coulomb cutoff of 1.0 nm was applied, and van der Waals interactions were smoothly switched to zero between 0.8 and 1.0 nm using a force-switch scheme. The temperature was maintained at 310 K using the v-rescale thermostat, and the pressure was maintained at 1 bar using the C-rescale barostat with isotropic coupling. Trajectory stability was evaluated using the RMSD and Rg of the actin trimer. Unless otherwise specified, quantitative interfacial and conformational analyses were performed over the final 20 ns of each production trajectory.

### Interfacial interaction analysis

Interfacial interactions within the actin–VASP complex were characterized by the buried solvent-accessible surface area (BSA), MM/GBSA binding energy, hydrogen bonds, and salt bridges. For an interface formed by components A and B, the BSA was calculated as

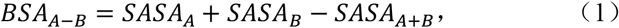

where SASA_*A*_and SASA_*B*_ are the solvent-accessible surface areas of components A and B, respectively, and SASA_*A*+*B*_is the solvent-accessible surface area of the corresponding complex. No factor of 1/2 was applied; therefore, the reported BSA represents the total solvent-accessible surface area buried on both sides of the interface (34–36). For all-atom trajectories,

SASA was calculated using a probe radius of 0.14 nm. Unless otherwise specified, BSA analyses were performed over the final 20 ns of each all-atom production trajectory. For coarse-grained trajectories, to characterize the long-timescale evolution of internal interfaces within the actin trimer, BSA was additionally calculated. Bead types were obtained directly from the corresponding topology (.itp) files. Regular, small, and tiny Martini 3 beads were assigned radii of 0.264, 0.230, and 0.191 nm, respectively. These values correspond to bead-size-dependent Lennard-Jones contact radii and were used to account explicitly for the different dimensions of Martini 3 bead classes(12, 37, 38).

Interfacial energetics were estimated using MM/GBSA analysis for all-atom trajectories (39). The calculated energy comprised van der Waals, electrostatic, polar solvation, and nonpolar solvation contributions. Calculations were performed over the final 20 ns of each all-atom trajectory. Mean values and corresponding measures of dispersion were calculated across the analyzed trajectories. Per-residue energy decomposition was further performed to identify residues contributing most favorably to the corresponding interfaces, and the 20 residues with the most favorable energetic contributions were retained for subsequent analysis. MM/GBSA energies were used for relative energetic comparisons among interfaces and conditions and were not interpreted as absolute binding free energies.

Interfacial hydrogen bonds were analyzed using the HydrogenBondAnalysis module of MDAnalysis 2.10.0 with Python 3.13.11. Donor, hydrogen, and acceptor atoms were identified according to the CHARMM36 topology (40, 41). A hydrogen bond was defined by a donor–acceptor distance of ≤3.5 Å and a D–H···A angle of ≥150°. For each residue pair, hydrogen-bond occupancy was calculated as

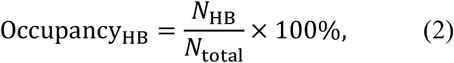

where *N*_HB_is the number of analyzed frames in which a given residue pair formed at least one hydrogen bond and *N*_total_is the total number of analyzed frames. Multiple atom-level hydrogen bonds between the same residue pair within a single frame were counted as one residue-pair interaction for occupancy calculations.

Salt bridges were analyzed using the distance-analysis routines implemented in MDAnalysis 2.10.0 with Python 3.13.11. A salt bridge was defined when oppositely charged side-chain atoms were separated by ≤4.0 Å (40, 41). Salt bridges were treated as nondirectional interactions, and each residue pair was counted at most once per frame regardless of the number of atom pairs satisfying the distance criterion. Salt-bridge occupancy was calculated as

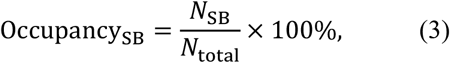

where *N*_SB_is the number of analyzed frames in which a given residue pair formed at least one salt bridge and *N*_total_is the total number of analyzed frames. Periodic boundary conditions were accounted for in all distance-based analyses.

### Actin conformational analysis and statistical analysis

To characterize subunit-specific conformational states within the actin trimer, three structural descriptors were analyzed for actin subunits A1, A2, and A3: W-loop gate distance, SD1–SD4 dihedral angle, and Asp239 displacement relative to an F-actin reference structure. Unless otherwise specified, conformational analyses were performed over the final 20 ns of each all-atom production trajectory, consistent with the interval used for interfacial analyses.

The W-loop gate state was characterized by the distance between the side-chain hydroxyl oxygen of Tyr138 and the backbone carbonyl oxygen of Gly163 within each actin subunit. This distance reports the opening and closure of the W-loop gate. Shorter distances correspond to a closed gate, whereas larger distances indicate an open gate (7). Conformations with distances below 7.5Å were classified as closed, whereas those with distances at or above this threshold were classified as open. The fractional occupancies of the closed and open states were calculated separately for A1, A2, and A3 over the analysis interval.

To characterize global subdomain twisting, the SD1–SD4 dihedral angle was calculated for each actin subunit. The four actin subdomains were defined using the following residue ranges: SD1 (a.a 3-27,69-131,342-358); SD2 (a.a 30-35,49-64); SD3 (a.a 145-175,270-327); and SD4 (a.a 180-251) (4). These definitions were adapted from the actin subdomain definitions used in previous structural analyses, with residue indices mapped to the numbering of the present simulation system (4, 6). For each trajectory frame, the geometric center of the Cα atoms within each subdomain was calculated. The SD1–SD4 dihedral angle was then defined by the four subdomain centers in the order SD1–SD2–SD3–SD4. The resulting signed dihedral angle was expressed in the same angular convention used for the reference actin structures. Because dihedral angles are periodic variables, circular statistics were used to calculate mean orientations and angular dispersion.

Asp239 displacement relative to the F-actin reference conformation was used to characterize the positional deviation of the SD4-associated region. The barbed-end F-actin structure (PDB ID: 8F8R) was used as the reference (20). For each trajectory frame, each simulated actin subunit was aligned to the reference by least-squares fitting of the Cα atoms belonging to SD2 and SD3. Following alignment, the three-dimensional Euclidean distance between the Cα atom of Asp239 in the simulated structure and that of the corresponding residue in the reference structure was calculated. Smaller displacement values indicate closer positional correspondence of the SD4-associated region to the F-actin reference geometry (4).

## Discussion

This study investigated how VASP stabilizes an early actin nucleation by combining Martini 3 coarse-grained simulations with all-atom molecular dynamics simulations. Our results support a model in which VASP stabilizes the actin trimer through a spatially selective and multivalent interaction mechanism. Rather than interacting uniformly with all three actin subunits, VASP preferentially engages the laterally associated A2 subunit and is coupled to reinforcement of the A2– A1 and A2–A3 internal interfaces. The preferential interaction of VASP with A2 was consistently observed across all three independent replicate simulations (Figure S3), supporting this interaction pattern as a reproducible feature of the VASP–Actin system. This preference is mechanistically important because the laterally associated subunit A2 represents a particularly vulnerable component of the trimeric nucleus (3, 4). VASP binding therefore stabilizes the trimer by anchoring A2 and strengthening its contacts with the longitudinal actin dimer. This stabilization does not simply rigidify the trimer or drive all three actin subunits toward a common filament-like conformation. Instead, A1, A2, and A3 exhibit distinct local and global conformational responses to VASP association. These findings suggest that VASP stabilizes a nucleation-like actin trimer while preserving the subunit-specific conformational heterogeneity of the early assembly.

Actin nucleation represents a major kinetic barrier to filament assembly because early oligomeric intermediates are thermodynamically and kinetically unfavorable and are prone to dissociation (2, 3). Formation of a trimer is generally considered a critical step toward establishment of a productive actin assembly (2, 3). Structural analysis of a trimeric actin nucleus further supports an asymmetric architecture in which a laterally associated third subunit joins a longitudinal actin dimer (3, 4). Our simulations suggest that VASP may contribute to nucleation by increasing the persistence of such an already formed trimeric intermediate. Importantly, the present simulations did not model the spontaneous association of freely diffusing G-actin monomers and therefore do not establish that VASP directly catalyzes the initial monomer-association steps. Instead, the mechanism described here should be interpreted specifically as stabilization and remodeling of a preformed nucleation trimer. This distinction is important because stabilization of a short-lived intermediate can promote the subsequent probability of productive elongation.

This mechanism has an informative parallel with the recently resolved villin-stabilized actin nucleus. In that structure, villin contacts multiple actin subunits, and the V3V4 linker and headpiece region contribute to stabilization of A2 interactions with both A1 and A3 (4). These observations suggest a broader mechanistic principle in which actin nucleators can stabilize a trimeric nucleus by reinforcing interactions involving the vulnerable lateral component. The molecular implementations, however, are distinct. Villin achieves this stabilization through an organized multidomain architecture surrounding the actin nucleus, whereas tetrameric VASP in our simulations engages the trimer through a distributed and dynamically maintained interaction network. This distinction suggests that reinforcement of lateral-subunit interfaces does not require a single stereotyped nucleator architecture and may instead be achieved through structurally diverse actin-binding proteins. More broadly, other actin nucleators have also been proposed to promote nucleation by organizing or stabilizing otherwise unfavorable inter-subunit contacts within oligomeric actin assemblies (16, 17), supporting the idea that stabilization of vulnerable interfaces may represent a recurrent strategy for overcoming the actin nucleation barrier.

In this context, the effect of VASP may be described as conformational stabilization without conformational homogenization. By limiting destabilizing inter-subunit rearrangements while allowing distinct subunit-specific conformational states to persist, VASP may increase the lifetime of the nucleation trimer without constraining it to a single filament-like structural state. Such conformational heterogeneity may itself be an intrinsic feature of an early actin nucleus. Indeed, a previous study showed that the villin-stabilized trimer contains protomers occupying markedly different positions along the G-to-F conformational landscape (4), further indicating that formation of a stable trimeric nucleus does not require simultaneous conversion of all constituent subunits into a canonical F-actin state. Instead, the nucleus appears capable of accommodating an asymmetric ensemble in which individual subunits adopt conformations suited to their local interaction environments.

Several limitations should be considered when interpreting these findings. First, because no experimentally determined structure of the VASP tetramer is currently available, we used a model-derived structure. This introduces additional uncertainty, particularly for flexible and intrinsically disordered regions. Second, the coarse-grained simulations required elastic-network restraints, which may limit large intramolecular conformational rearrangements (25). Conclusions regarding local and subdomain-level conformational changes were derived primarily from the all-atom simulations. However, the all-atom simulaitons did not reach microsecond timescales and may consequently have failed to capture slow transitions between distinct conformational states of the actin subunits.Finally, because the simulations were initiated from a preassembled actin trimer, they cannot resolve the kinetics or pathways of spontaneous trimer formation. Future free-association simulations, enhanced-sampling approaches, and kinetic studies will be required to determine whether VASP also influences the preceding monomer-association steps and how stabilization of the trimer affects subsequent filament elongation.

Together, our findings support a molecular model in which VASP increases the persistence of a nucleation actin trimer by preferentially engaging the lateral A2 subunit and reinforcing its interactions with the longitudinal A1/A3 core, while preserving subunit-specific conformational heterogeneity. Rather than establishing an entirely new actin–actin architecture, VASP appears to reinforce an existing but intrinsically unstable intermediate. Our results provide a potential molecular basis for the previously reported nucleation-promoting activity of VASP (8, 18, 29) and extends the established view of VASP as an actin-binding protein that recruits monomers and promotes filament elongation. More broadly, the simulations suggest that actin nucleation may be facilitated not only by generating new inter-subunit contacts, but also by selectively stabilizing vulnerable interfaces while retaining the conformational asymmetry. Our results are derived from molecular simulations, but can be validated experimentally. The predicted A2-centered interaction network and high-contribution residue pairs, including Asp20(A2)–Lys239(VASP), Asp21(A2)–Lys239(VASP), and Glu329(A2)–Lys233(VASP), provide experimentally testable candidates for site-directed mutagenesis, actin polymerization assays, cross-linking approaches, or structural characterization. Experimental perturbation of the A2–VASP interaction network and longer-timescale simulations will be important for testing this model and determining how stabilization of the trimer is coupled to subsequent actin assembly.

## Acknowledgements

This research was funded by the National Natural Science Foundation of China 12404241, the Guangdong High-Level Talent Program 2024QN11X306, and the GJYC Program of Guangzhou 2025Y01Q0037.

