## Supplemental Figures for "VASP association stabilizes the actin nucleus revealed by molecular dynamics simulations"

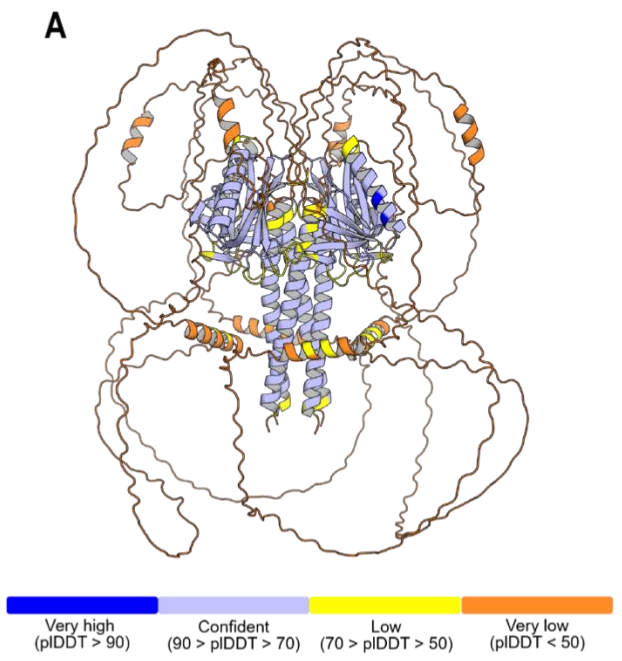


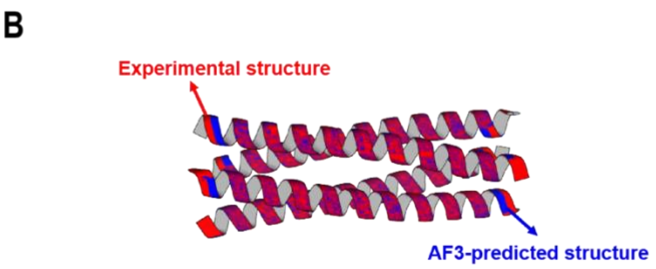


C

| Structural metric | Result |
|---|---|
| Sequence identity | 100% |
| Matched residues | 40 × 4 chains |
| Single-chain Cα RMSD (Å) | 1.63–1.66 |
| Tetramer Cα RMSD (Å) | 1.71 |
| Contact Jaccard index | 0.688 |
| Total SASA loss (nm²) | 70.71 vs 69.14 |

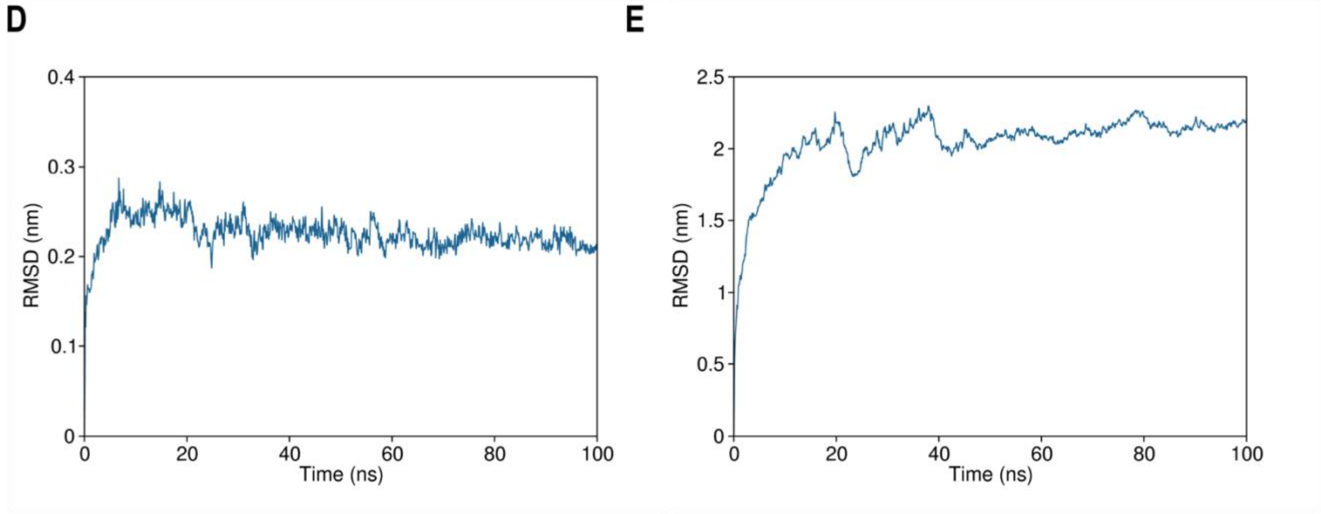


**Figure S1. Structural model and dynamic stability of the VASP tetramer during all-atom molecular dynamics simulation.** (A) Representative AlphaFold3-predicted full-length VASP tetramer, with the C-terminal tetramerization domain (residues 338–377) highlighted in different colors for the four VASP chains and the remaining regions shown in gray. Across 24 independent AlphaFold3 predictions, the tetramerization domain showed a mean pLDDT of 73.95 ± 1.57 and a mean inter-chain PAE of 8.32 ± 0.84 Å. Structural comparison among the 24 predictions (276 pairwise comparisons) yielded a permutation-optimized pairwise Cα RMSD of 0.541 ± 0.219 Å, indicating highly consistent convergence of the predicted tetramerization-domain architecture across independent predictions. (B) Structural alignment of the predicted VASP tetramerization domain (blue) with the experimentally determined tetrameric domain (orange). (C) Quantitative comparison between the predicted VASP tetramerization domain and the experimentally resolved tetrameric domain. (D) Cα root-mean-square deviation (RMSD) of the tetramerization domain during the 100-ns all-atom simulation. Following a short initial conformational adjustment, the tetramerization core remained relatively stable, with RMSD values fluctuating mainly around 0.2–0.25 nm. (E) RMSD of the full-length VASP tetramer during the same simulation. The full-length structure underwent substantial conformational rearrangement during the early stage of the simulation and subsequently reached a relatively stable plateau at approximately 2.0–2.2 nm. Together, these results indicate that the predicted full-length VASP tetramer is structurally stable throughout the simulation.

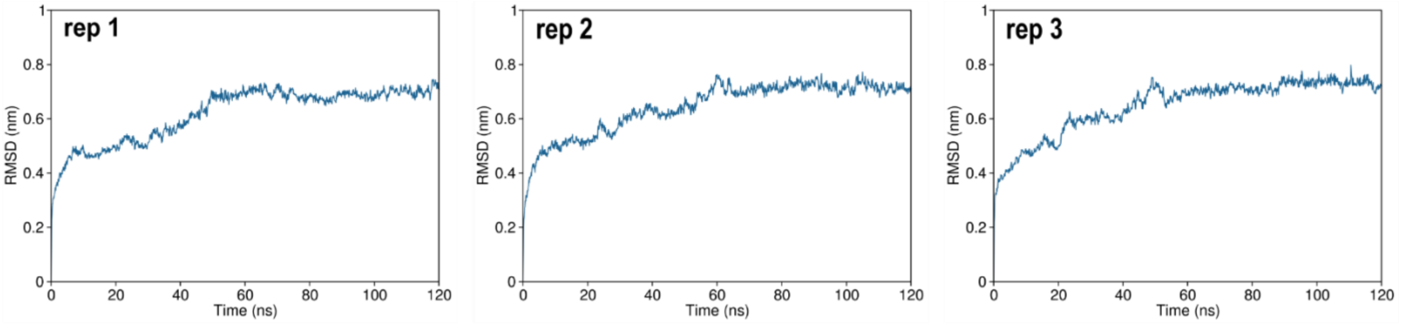


**Figure S2. RMSD profiles of the Actin trimer after backmapping to the all-atom representation.** The backbone root-mean-square deviation (RMSD) of the Actin trimer was monitored over three independent 120-ns all-atom molecular dynamics simulations following backmapping from the coarse-grained model. In all replicates, the RMSD increased during the early stage of the simulations and subsequently fluctuated around a relatively stable level of approximately 0.7 nm, indicating equilibration of the Actin trimer during the later stage of the all-atom simulations ( > 60 ns).

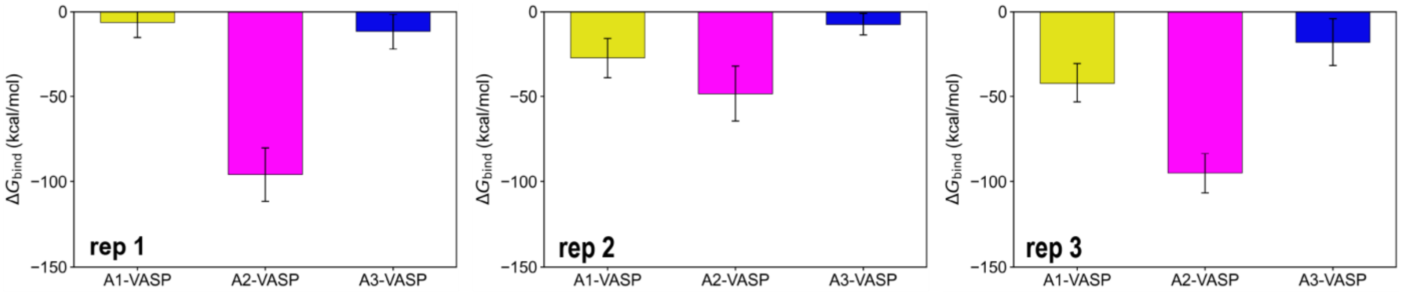


**Figure S3. Subunit-specific VASP binding free energies for the Actin trimer across three independent simulations.** The MM/GBSA-derived binding free energy contributions $\Delta G(_{\mathrm{bind}})$ between VASP and each Actin subunit (A1, A2, and A3) are shown for three independent replicates (rep 1–3, left to right). Bars represent the mean $\Delta G(_{\mathrm{bind}})$ values for each replicate, and error bars indicate the corresponding standard deviation within that replicate. More negative values indicate more favorable VASP–Actin interactions. Across the three replicates, A2 consistently exhibited the most favorable interaction with VASP, whereas A1 and A3 showed weaker binding contributions.
